# Terrestrial DNA viromes are systematically more divergent from reference databases than marine ones, with polar ecosystems amplifying the contrast

**DOI:** 10.64898/2025.12.09.693195

**Authors:** Vaibhav Kulkarni, Stéphane Aris-Brosou

**Author notes:** Corresponding author: Department of Biology, University of Ottawa, Ottawa, ON K1N 6N5, Canada.

## Abstract

Viruses are the most abundant biological entities on Earth, yet most of their diversity remains unsampled and unevenly represented across ecosystems. To test whether viral novelty is structured more by biome or latitude, we analyzed 120 curated environmental DNA metagenomes spanning marine and terrestrial habitats across equatorial temperate, Arctic, and Antarctic regions. Using a unified metagenomic and phylogenomic workflow, we reconstructed viral protein families, linked environmental sequences to the IMGVR v4.1 reference collection, and quantified divergence using three complementary measures: within-clade patristic distance, the bridging edge to the nearest reference neighbor, and phylogenetic tree-shape statistics. Because distances within phylogenetic trees are not independent, all inference was performed at the sample level. Across all samples, reconstructed environmental viral sequences were substantially more divergent from current reference genomes than their database homologs. The dominant pattern was biome-driven: terrestrial viromes were consistently farther from IMGVR references than marine viromes in every latitudinal band, and this contrast was supported by both branch-length and topology-based metrics. Latitude modulated this baseline signal, with polar ecosystems, especially Antarctic terrestrial samples, occupying the most divergent end of the two-axis landscape. Tree imbalance statistics showed that terrestrial viral phylogenies were more deeply branching and more asymmetric than marine ones, providing independent support for the biome effect. In contrast, some previously reported polar contrasts weakened after correcting for pseudoreplication and using proper sample-level replication. These results argue that viral novelty is structured primarily by biome, with polar environments amplifying an already strong terrestrial–marine contrast. The findings also indicate that current reference databases remain disproportionately sparse for terrestrial, especially polar terrestrial, viromes. Expanding viral sampling in these environments will be essential for improving ecological inference, improving database coverage, and refining our understanding of global viral diversity.

## Introduction

Viruses are the most abundant biological entities on Earth, outnumbering cellular life by at least an order of magnitude and comprising an estimated 10^31^ particles globally [1, 2]. They are key regulators of microbial evolution and ecosystem function, mediating horizontal gene transfer and modulating elemental cycling across marine, freshwater, and terrestrial biomes [3, 4, 5]. Viral infection-driven lysis, the viral shunt, reallocates organic carbon and nutrients, restructuring food webs and influencing biogeochemical feedbacks linked to climate processes. Through high rates of mutation, recombination, and reas-sortment, viruses constitute a vast and dynamic gene pool that shapes host adaptability and ecosystem resilience [6, 7].

Despite their ubiquity and ecological significance, viral genomes remain the least characterized component of global biodiversity. More than 99% of viral genetic diversity, often referred to as viral dark matter, is uncatalogued, limiting ecological inference and predictive modeling [8]. Large-scale initiatives such as the Global Virome Project [9] and the IMG/VR database [10] have expanded viral reference collections, yet substantial gaps persist. More critically, these gaps are not evenly distributed across global ecosystems: reference databases have historically been dominated by viromes from marine ecosystems (most prominently the Tara Oceans datasets and related ocean surveys [11]), while terrestrial viromes, particularly from soils, remained comparatively undersampled until the IMG/VR v4 release [12, 10]. Independent of latitude, this differential reference coverage predicts that environmental queries from terrestrial sources will recruit more distant IMG/VR neighbors than marine queries.

This prediction of a biome-level asymmetry is reinforced by underlying microbial-ecology principles. Soils are spatially structured at the millimeter scale, with pH, water potential, organic-matter quality, and oxygen gradients producing vastly more independent ecological niches per unit volume than seawater [13]. Soil microbial communities are correspondingly more diverse and more dispersal-limited than marine ones, and the viral component of these communities inherits this structure through tight host coupling [12, 14]. Soils also archive ancient microbial and viral lineages through permafrost preservation, with viable giant viruses recovered from Siberian permafrost demonstrating that some terrestrial polar viromes integrate evolutionary signal across millennial timescales [15].

Latitude provides a complementary axis. Classical ecological theory predicts a latitudinal diversity gradient (LDG), with species richness decreasing toward the poles [16, 17], and several recent metagenomic surveys have nonetheless found polar Arctic and Antarctic ecosystems to be unexpected hotspots of viral novelty [18, 11, 19, 20, 12, 21]. The Arctic has warmed nearly four times faster than the global mean since 1979 [22], and polar regions impose strong physicochemical gradients, extreme seasonality, and oligotrophic conditions that drive idiosyncratic viral diversification [23, 24, 25], all amplified by sea ice loss [26], altered hydrological cycles [27], and permafrost thaw [28]. Existing polar virome studies, however, have been fragmented by methodological inconsistencies and uneven geographic coverage [29, 30, 31], and they have rarely been placed alongside a comparably-processed temperate baseline. Whether polar viromes are special in absolute terms, or whether what looks like a polar effect is in part a biome effect propagated through differentially represented latitudes, has therefore not been resolved.

Here we present a latitudinally balanced comparison of environmental DNA viromes drawn from marine and terrestrial ecosystems across equatorial / temperate, Arctic, and Antarctic regions. Using a unified analytical workflow, we reconstruct viral genes from metagenomic data and quantify three complementary signatures of divergence from the IMG/VR v4.1 reference collection: within-clade pairwise patristic distances among reconstructed sequences (‘gene mpd’), the bridging edge connecting the reconstructed clade to its closest IMG/VR neighbor, and three tree-shape statistics that capture the topology of the recovered phylogenies (Pybus-Harvey *γ*, Yule-normalized Colless and Sackin imbalance). We then test, at the proper sample-level resolution, whether the polar-versus-temperate contrast is statistically supported and, importantly, whether a marine-versus-terrestrial contrast operates on the same data. We find that the biome contrast is the larger and more consistent of the two: terrestrial viromes are systematically more divergent from current references than marine ones across all latitudinal bands, with polar ecosystems amplifying the contrast.

## Materials and Methods

### Study design and sample selection

An overview of the analytical workflow is provided in Fig. 1. A total of 180 publicly available environmental DNA metagenomes were retrieved from the NCBI Sequence Read Archive (SRA), stratified across two biomes (terrestrial, marine) and three latitudinal zones (north polar >66.5°N, equatorial / temperate 66.5°N-66.5°S, and south polar >66.5°S), with 30 metagenomes per biome / latitude combination. We call each of these combination a *stratum*. Geographic spread was maximized under the constraint of existing SRA data within each stratum. After all quality control steps, 120 datasets met inclusion criteria.

**Figure 1.**
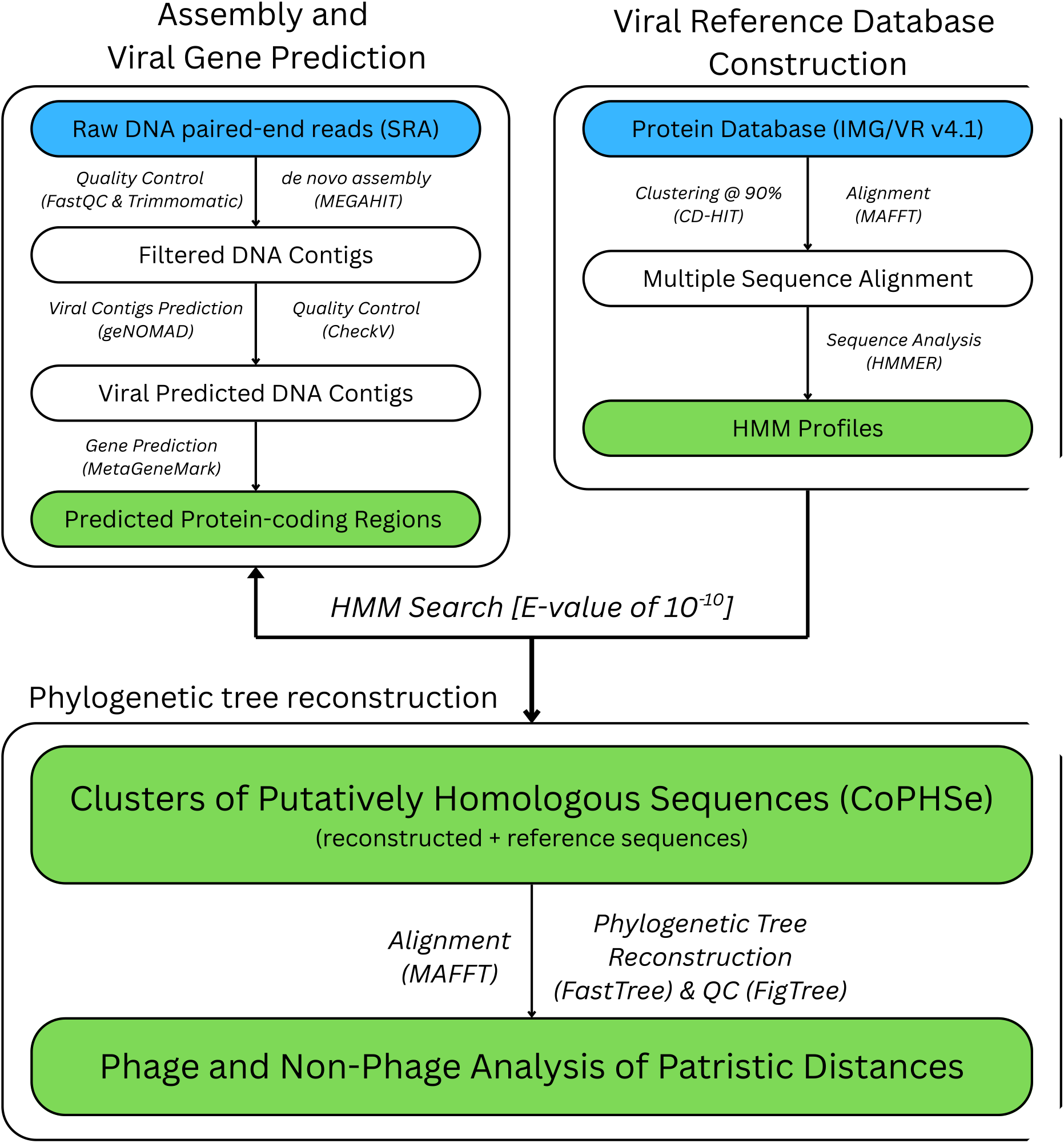
Viral metagenomic analysis pipeline. Raw DNA paired-end reads (SRA) were assembled and protein-coding regions predicted. Together with IMG/VR v4.1 reference proteins, HMM profiles were built to generate clusters of putatively homologous sequences (CoPHSe). Patristic distances and tree-shape statistics were then computed for phage and non-phage groups.

### Read processing and *de novo* assembly

Raw sequencing reads were downloaded using the SRA Toolkit v3.0.0 and evaluated with FastQC v0.12.1 [32, 33]. Adapter trimming and quality filtering were performed with Trimmomatic v0.39 [34]. High-quality reads were assembled *de novo* using MEGAHIT v1.2.9 under default settings [35]. Assembly contiguity (N50) was tracked alongside the divergence metrics in the sample-level analyses below (Fig. S1, S2).

### Viral contig identification and quality assessment

Assembled contigs were processed with geNomad v1.11.1 [36] and screened with CheckV v1.0.3 [37]; only medium- to high-quality and complete contigs were retained. Protein-coding sequences were predicted using MetaGeneMark v3.38 [38].

### Reference database

Uncultivated viral reference sequences were obtained from IMG/VR v4.1 (release 2022-09-20) [10]. The IMG/VR protein dataset was dereplicated at 90% sequence identity using CD-HIT v4.8.1 [39, 40]. Host and taxonomic annotations were used to categorize phylogenetic trees as phage, non-phage, or unknown.

### Homology clustering and HMM profile construction

Dereplicated IMG/VR proteins were clustered and aligned with MAFFT v7.471 [41, 42]. Alignments containing ≥ 10 sequences were converted into profile Hidden Markov Models (HMMs) using HMMER v3.2.1 [43]. Each HMM profile was searched against the predicted proteins from environmental contigs using an E-value cutoff of 10^−10^. Matches were grouped into Clusters of Putatively Homologous Sequences (CoPHSe; [21]) containing both environmental and IMG/VR proteins.

### Phylogenetic tree construction and filtering

Each CoPHSe was realigned with MAFFT and trimmed with trimAl v1.4 (`-gappyout`) [44]. Phylogenetic trees were inferred with FastTree v2.1.11 [45] under the LG substitution model. Most CoPHSe trees formed monophyletic environmental clades; non-monophyletic cases were excluded (Fig. S3). Trees were parsed in R using the APE package v5.8.1 [46]. Trees with more than 400 IMG/VR reference sequences were pruned to that threshold to bound the quadratic scaling of patristic distance calculations. Each tree was then subdivided into two subtrees: one consisting solely of reconstructed sequences (tagged “gene”) and one of reference sequences (tagged “IMG/VR”).

### Divergence metrics and statistical analysis

For each tree, we computed (i) the mean pairwise patristic distance (MPD) among re-constructed tips (gene_mpd), (ii) the MPD among IMG/VR reference tips (imgvr_mpd), and (iii) the bridging edge connecting the most recent common ancestor of each sub-group, which captures the path length from the reconstructed clade to its closest IMG/VR neighbor (Fig. S4). Because pairwise patristic distances within a CoPHSe tree are not independent observations, statistical inference was carried out at the sample level: each SRA accession was reduced to one observation per metric (the median across all of that sample’s CoPHSe), and Wilcoxon rank-sum tests with Cliff’s *δ* effect sizes and bootstrap 95% confidence intervals were used to compare biomes (marine versus terrestrial) and latitudinal bands (Arctic / Antarctic versus equatorial). Comparisons were performed both pooled across the orthogonal axes and within each level of the orthogonal axis.

### Tree-shape statistics

For every CoPHSe with at least six environmental tips, three independent tree-shape statistics were computed: the Pybus-Harvey *γ* statistic [47], the Yule-normalized Colless imbalance index [48], and the Yule-normalized Sackin index [49]. Trees were midpoint-rooted with `phangorn::midpoint` and resolved to fully bifurcating topologies with `ape::multi2di` prior to indexing. Sample-level Wilcoxon contrasts were performed on per-SRA medians, identically to the divergence metrics.

## Results

### Dataset quality control and filtering

A total of 180 environmental metagenomes were assembled with MEGAHIT v1.2.9. Regional N50 distributions were largely comparable across terrestrial strata (Kruskal-Wallis *H* = 5.12, *p* = 0.077); a modest difference was detected for marine strata (*H* = 6.54, *p* = 0.038). We explicitly accounted for this in the sample-level analyses below by tracking N50 alongside each per-SRA divergence value, and verified that the biome and latitude effects reported here persist across the full N50 range represented in the cohort (Figs. S1-S2). Seventeen assemblies (1 terrestrial, 16 marine) did not meet minimum QC criteria and were excluded. The final dataset consisted of 120 curated assemblies evenly distributed across terrestrial and marine biomes within the three latitudinal bands (Fig. 2; Tables S1-S2).

**Figure 2.**
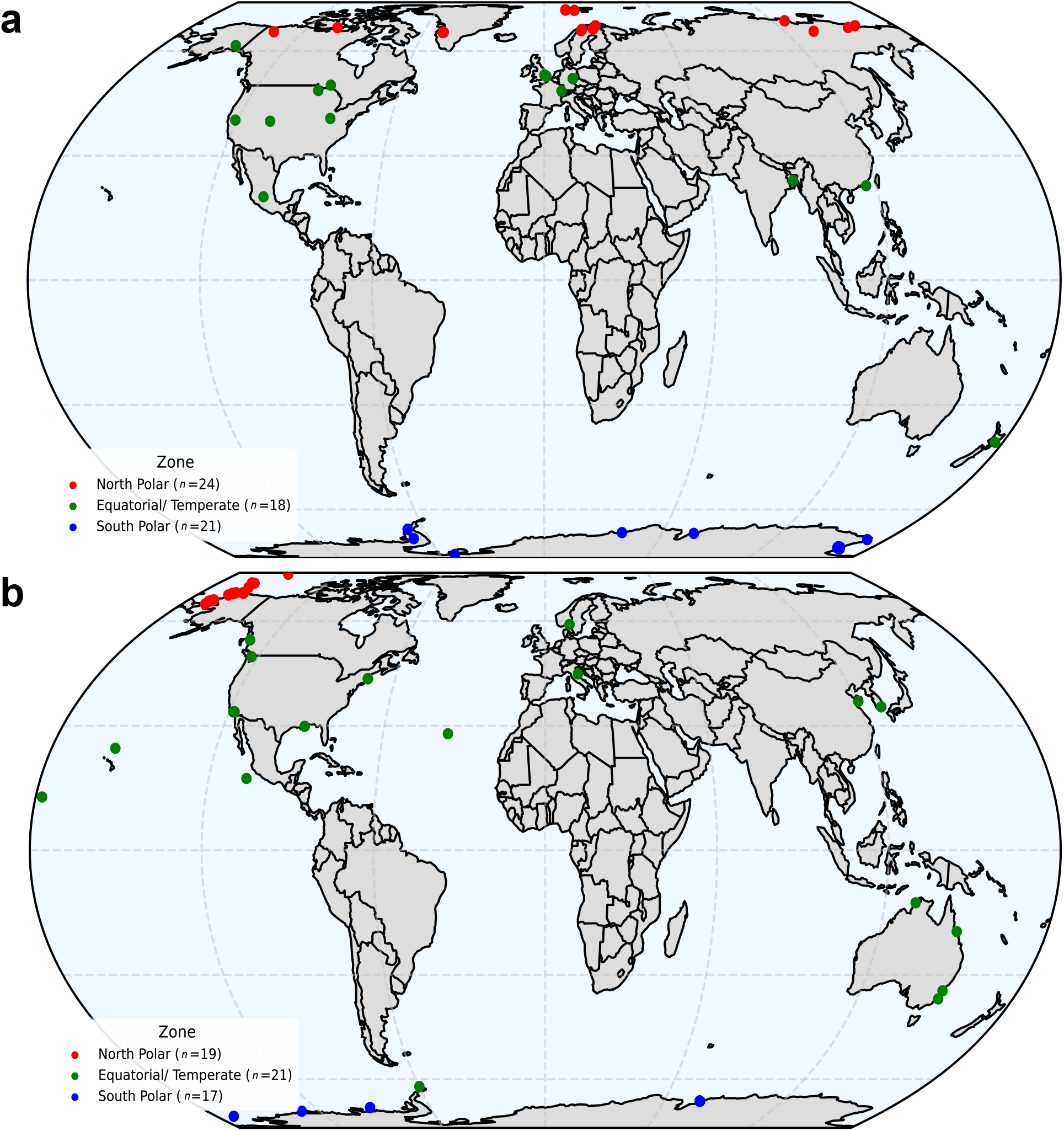
Global distribution of analyzed viral metagenomic samples. Equal-Earth projection maps illustrating the geographic locations of samples that passed assembly and quality control procedures. **(a)** Terrestrial (*n* = 63) and **(b)** marine samples (*n* = 57) are shown separately, highlighting the latitudinal coverage and spatial representation of the datasets used in downstream analyses.

### Global patterns of viral genetic divergence

From these 120 assemblies (17-24 per stratum), predicted viral proteins were clustered with their closest IMG/VR analogs to generate CoPHSe. A limit of 400 IMG/VR references per tree was applied (excluding 0.35% of cases; Fig. S5), yielding 1,846,393 valid phylogenies. For each tree, three complementary metrics were derived. Across all biomes and latitudes, reconstructed environmental sequences displayed markedly greater divergence than their database homologs (Table 1; Figs. 3-4). Median gene mpd values for reconstructed sequences ranged from 1.4 to 1.9 substitutions per site, bridging-edge medians from 0.6 to 0.9, and IMG/VR reference medians from 0.02 to 0.05, confirming that reconstructed environmental viruses occupy substantially broader phylogenetic space than existing reference databases capture.

**Table 1.** Median patristic distances (MPD) estimated across biomes, taxonomy and regions. Bridging-edge medians and corresponding sample-level Wilcoxon contrasts are reported in Supplementary Tables S3 and S4.

| Biome | Taxonomy | Region | MPD (subs/site) |  |  | Total trees (*n*) |
| --- | --- | --- | --- | --- | --- | --- |
|  |  |  | Bridging edge | Reconstructed sequences | IMG/VR sequences |  |
| Marine | Phage | North | 0.652 | 1.320 | 0.037 | 116,226 |
|  |  | Temperate | 0.655 | 1.458 | 0.032 | 710,703 |
|  |  | South | 0.710 | 1.912 | 0.030 | 454,700 |
|  | Non-phage | North | 1.530 | 0.126 | 0.024 | 190 |
|  |  | Temperate | 0.779 | 1.690 | 0.029 | 2,354 |
|  |  | South | 0.873 | 1.967 | 0.029 | 1,577 |
| Terrestrial | Phage | North | 0.844 | 2.176 | 0.027 | 175,182 |
|  |  | Temperate | 0.861 | 1.971 | 0.027 | 267,462 |
|  |  | South | 0.834 | 2.223 | 0.028 | 115,839 |
|  | Non-phage | North | 1.076 | 1.790 | 0.024 | 925 |
|  |  | Temperate | 1.231 | 1.578 | 0.026 | 587 |
|  |  | South | 0.834 | 2.270 | 0.033 | 648 |

**Figure 3.**
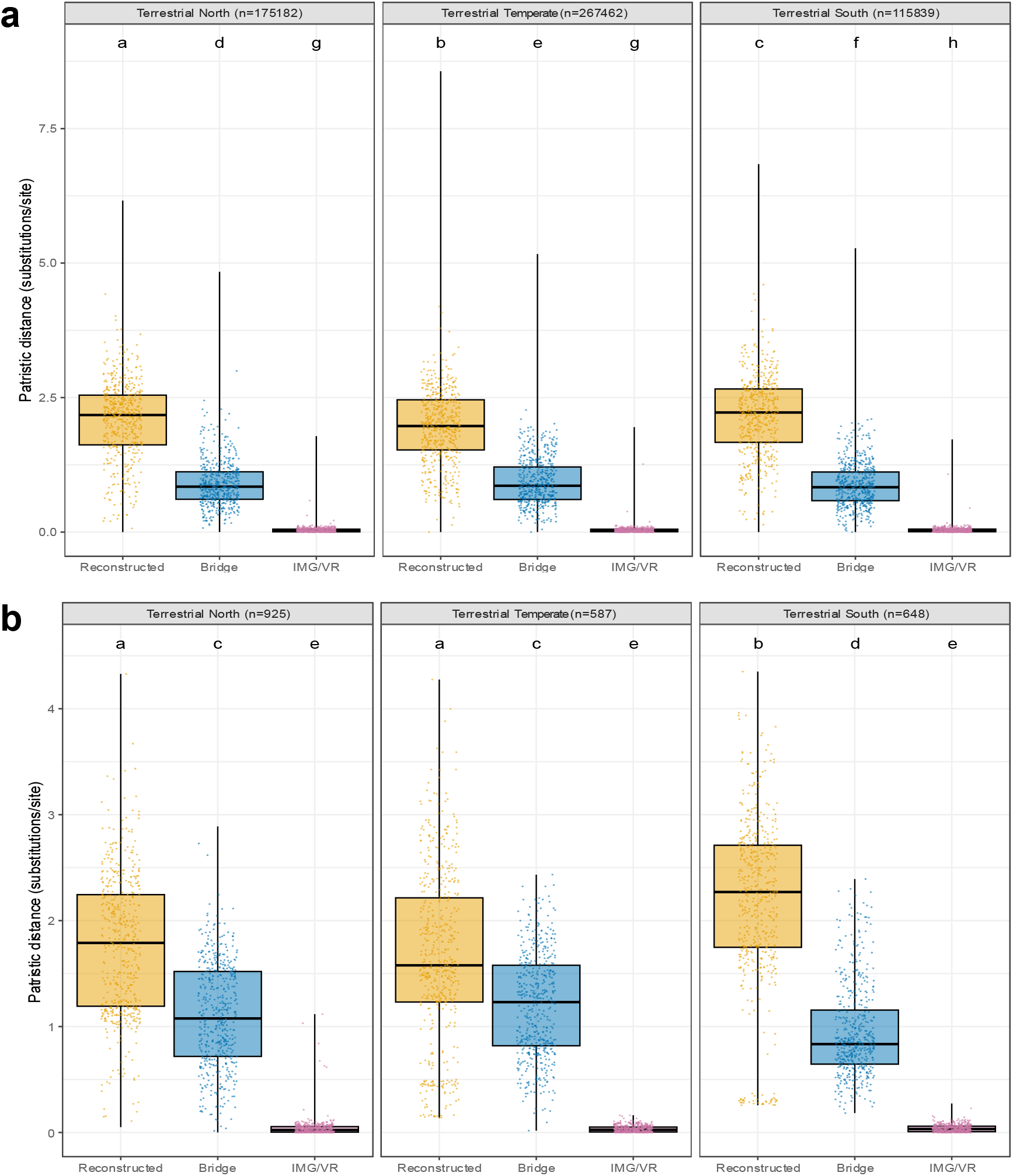
Patristic distance distributions in terrestrial viral communities. Boxplots show phylogenetic divergence for **(a)** phage and **(b)** non-phage sequences in terrestrial samples. Distances are derived from reconstructed environmental sequences compared with IMG/VR v4.1 references. Statistical significance shown here was originally assessed using Kruskal-Wallis tests on the full set of within-tree pairwise distances; because these distances are not independent observations, those *χ*^2^ values overstate the signal. Sample-level Wilcoxon contrasts using one observation per SRA are reported in Tables S3 (latitudinal contrasts) and S4 (biome contrasts).

**Figure 4.**
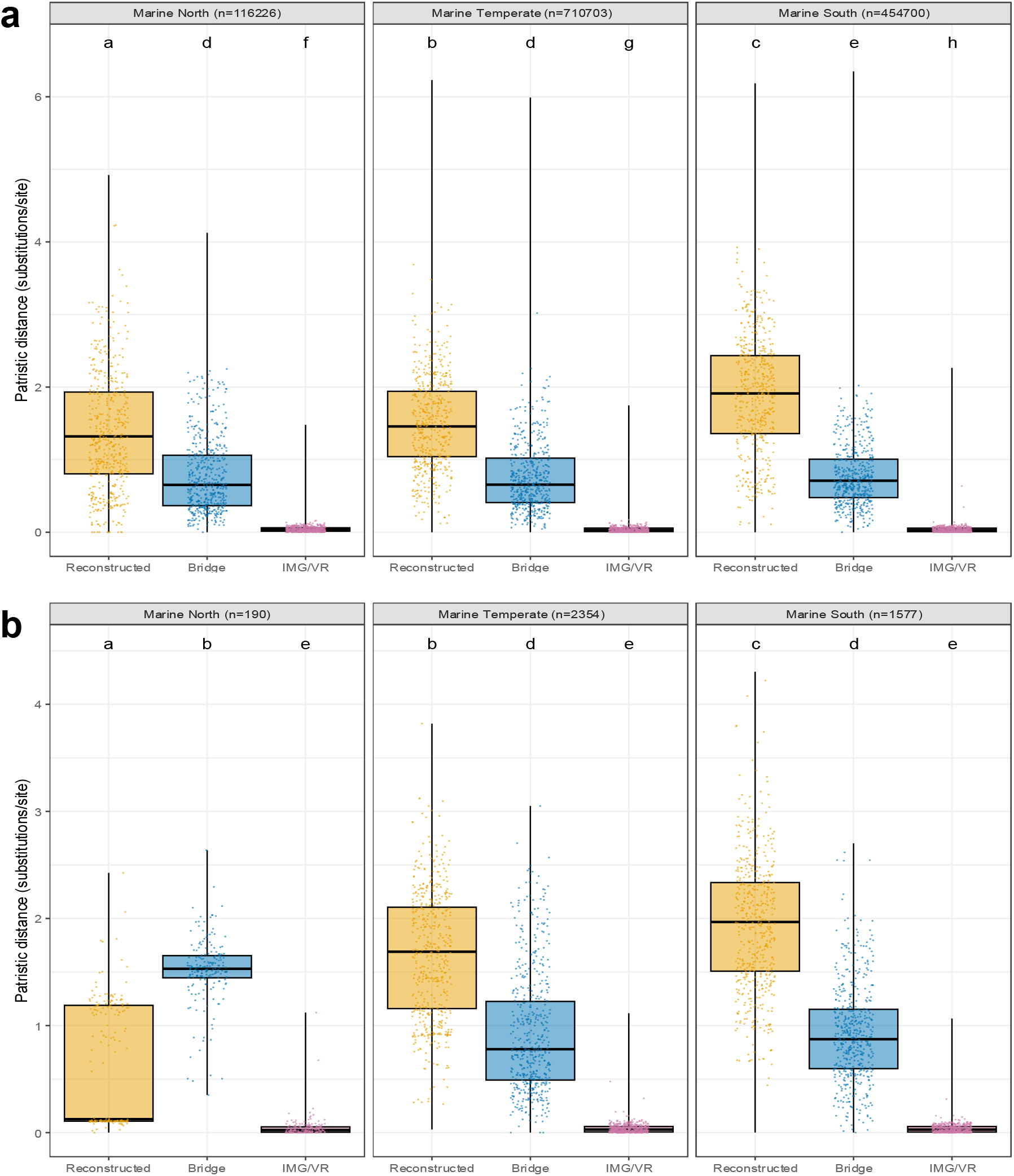
Patristic distance distributions in marine viral communities. Boxplots show phylogenetic divergence for **(a)** phage and **(b)** non-phage sequences in marine samples. Distances were calculated from reconstructed environmental sequences relative to IMG/VR v4.1 references. As in Fig. 3, the per-distance Kruskal-Wallis statistics shown here are inflated by pseudoreplication of within-tree pairwise distances; sample-level Wilcoxon tests are reported in Tables S3 and S4.

### Terrestrial viromes are systematically further from reference databases than marine viromes

At the proper sample-level resolution, the marine-versus-terrestrial contrast is the largest and most consistent signal in the data (Fig. 5). Per-SRA medians of the bridging-edge metric were significantly higher in terrestrial than in marine samples in every latitudinal band, with effect sizes (Cliff’s *δ*) ranging from moderate to large (equatorial / temperate *δ* = −0.63, *p* = 8 *×* 10^−4^, *n*_M_ = 21 [Marine] versus *n*_T_ = 18 [Terrestrial]; Arctic *δ* = −0.38, *p* = 0.037, *n*_M_ = 19 versus *n*_T_ = 24; Antarctic *δ* = −0.59, *p* = 0.002, *n*_M_ = 17 versus *n*_T_ = 21; Supplementary Table S2). Pooled across the three latitudinal bands, the biome contrast on bridging edge reached *p* < 10^−5^ with Cliff’s *δ* = −0.50. By contrast, within-clade gene mpd showed no significant biome contrast in any band, consistent with the limited sample-level power observed for the latitudinal contrasts on the same metric (see below). The within-IMG/VR reference MPD itself differed between biomes in a region-dependent way (equatorial: marine > terrestrial, *δ* = +0.59, *p* = 0.002; Antarctic: marine < terrestrial, *δ* = −0.52, *p* = 0.007; Fig. S6), reflecting the latitudinal composition of the IMG/VR collection itself. The biome contrast on bridging edge is approximately five times larger in magnitude than this reference-set asymmetry and therefore not explained by it. Full per-CoPHSe distributions of the bridging-edge metric across all 120 samples are shown in Fig. S7.

**Figure 5.**
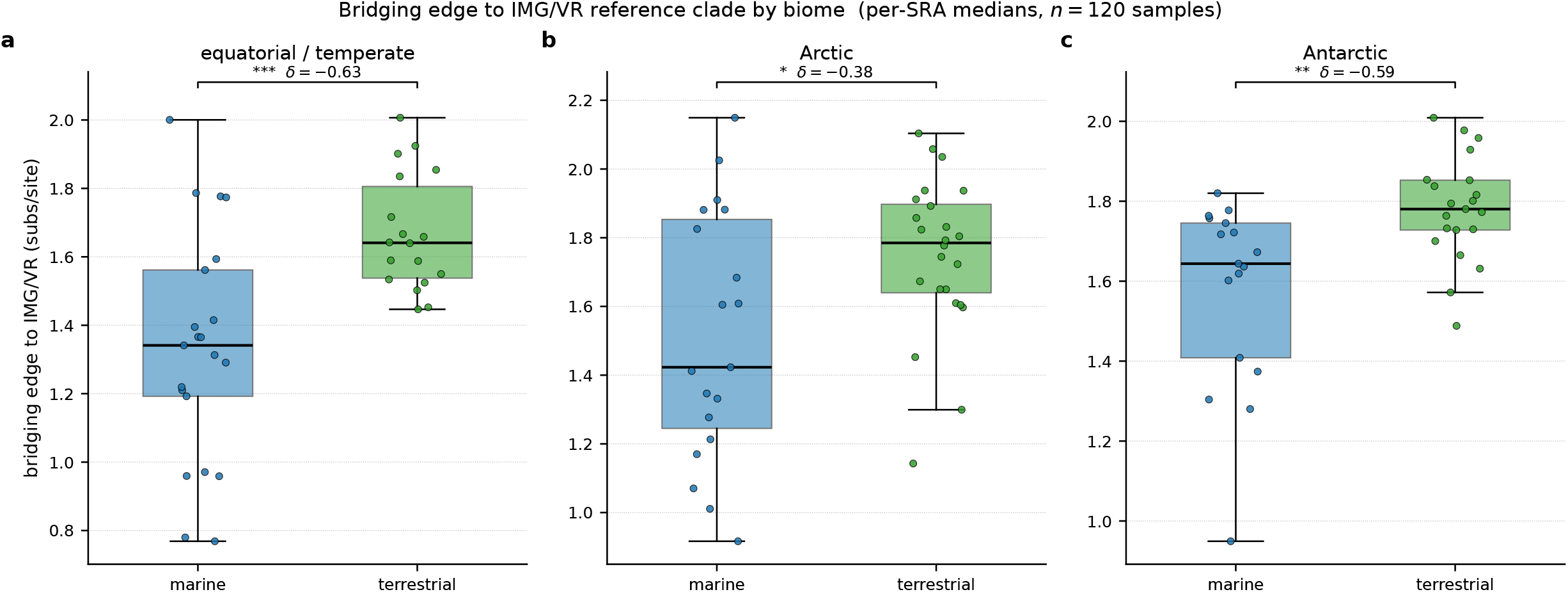
Terrestrial viromes are systematically further from IMG/VR reference clades than marine viromes in every latitudinal band. Boxplots and per-SRA jittered points of the bridging-edge metric, with biome on the x-axis and one sub-panel per latitudinal band (**a**, equatorial / temperate; **b**, Arctic; **c**, Antarctic). Each dot is one SRA accession (*n*=120 samples in total). Significance brackets show sample-level Wilcoxon rank-sum contrasts between marine and terrestrial within each band, with Cliff’s *δ* effect sizes inline (* *p* < 0.05, ** *p* < 0.01, *** *p* < 0.001). The biome contrast is significant in every latitudinal band and is consistently directional (terrestrial > marine), with effect sizes from *δ* = −0.38 (Arctic) to *δ* = −0.63 (equatorial). Corresponding biome-axis tree-shape contrasts are shown in Fig. S8 and the full 24-row contrast table is given in Table S4.

**Figure 6.**
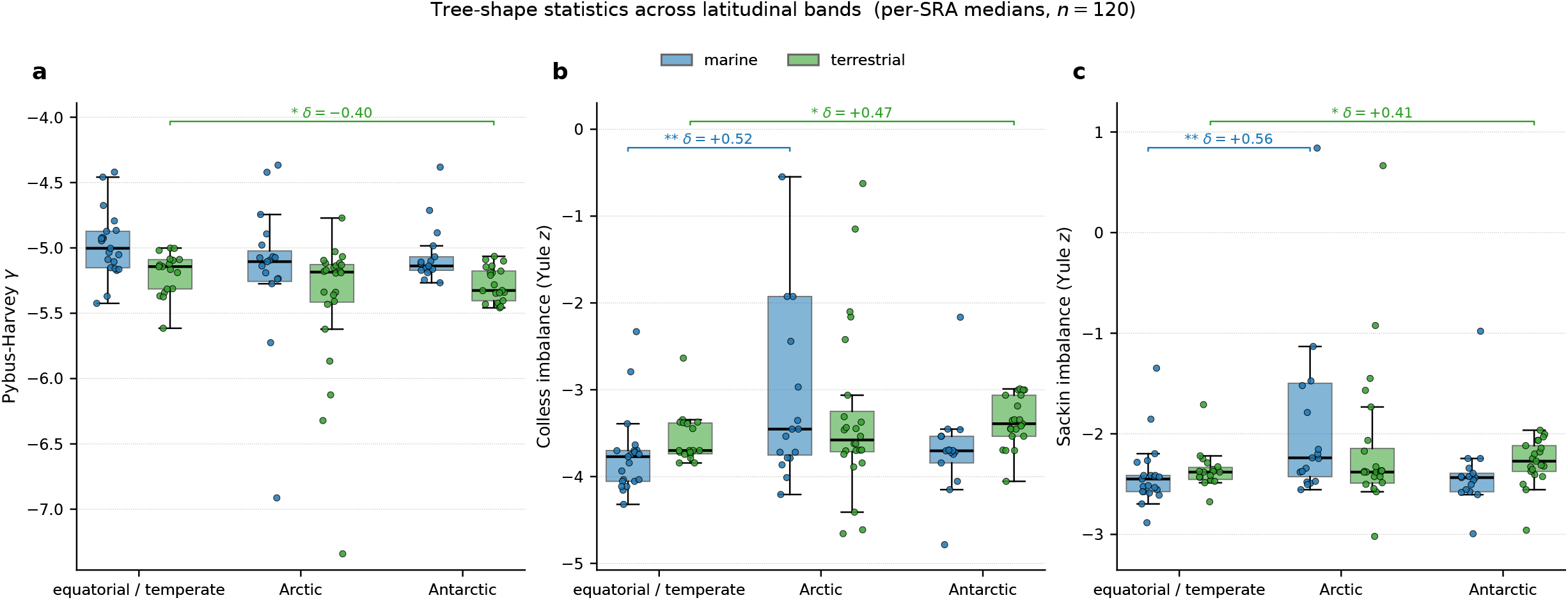
Tree-shape statistics across latitudinal bands at sample-level resolution. Boxplots and per-SRA jittered points for **(a)** the Pybus-Harvey *γ* statistic, **(b)** Yule-normalized Colless imbalance, and **(c)** Yule-normalized Sackin imbalance, computed on every CoPHSe with ≥ 6 environmental tips across all 120 samples (*n*_trees_ = 6,201,297). Significance brackets show sample-level Wilcoxon rank-sum contrasts versus the equatorial / temperate band of the same biome; only contrasts with *p* < 0.05 are annotated, with Cliff’s *δ* effect sizes inline. Marine north (panels b and c) and terrestrial south (all three panels) show significant elevation in tree imbalance and excess of early branching relative to equatorial samples. The complementary biome-axis view is shown in Fig. S8.

### Tree-shape statistics confirm the biome effect

Three independent tree-shape statistics were computed across all 120 samples (6,201,297 trees with ≥ 6 tips). Sample-level Wilcoxon contrasts confirmed that terrestrial trees are topologically more deeply branching and more imbalanced than marine ones, with the strongest effects in the equatorial and Antarctic bands (Fig. S8; Pybus-Harvey *γ* pooled *δ* = +0.43, *p* < 10^−4^, terrestrial trees more negative; Colless imbalance pooled *δ* = −0.27, *p* = 0.010, terrestrial trees more imbalanced). Within the Antarctic band, all three tree-shape statistics showed significant biome contrasts (*γ*: *δ* = +0.56, *p* = 0.004; Colless: *δ* = −0.67, *p* = 4 *×* 10^−4^; Sackin: *δ* = −0.56, *p* = 0.004). Within the equatorial band, *γ* and Colless were significant; Arctic contrasts trended in the same direction without reaching *α* = 0.05. Because tree-shape statistics depend on branching topology rather than pairwise branch lengths, they provide an orthogonal line of evidence for the biome effect. Full per-CoPHSe distributions of the three tree-shape statistics across all 120 samples are shown in Fig. S9.

### Polar ecosystems amplify the biome contrast

Although the biome effect is the dominant signal, polar ecosystems further modulate divergence. At sample-level resolution, the bridging-edge contrast between south polar and equatorial samples reaches *p* < 0.05 in both biomes (marine Δ = +0.30 subs/site, *δ* = +0.43, *p* = 0.026; terrestrial Δ = +0.14, *δ* = +0.41, *p* = 0.029; Table S3). The corresponding north polar contrasts trend in the same direction (marine *δ* = +0.28, *p* = 0.14; terrestrial *δ* = +0.31, *p* = 0.09) without reaching significance. The within-clade gene mpd contrast is directionally consistent (marine south Δ = +0.49, *δ* = +0.29, *p* = 0.14; terrestrial south Δ = +0.28, *δ* = +0.19, *p* = 0.34) but underpowered at the current sample count, an outcome attributable to the small effective sample size (*n*=16-24 per stratum) given the moderate effect. The intraclass correlation coefficient on per-CoPHSe gene mpd is 0.235, indicating that 76.5% of variance lies within rather than between samples and explaining why the original per-distance Kruskal-Wallis *χ*^2^ statistics dramatically overstate the latitudinal signal. Tree-shape contrasts identify additional polar-specific signatures: south polar terrestrial samples have significantly more negative *γ* (*δ* = −0.40, *p* = 0.036) and more imbalanced Colless (*δ* = +0.47, *p* = 0.014) and Sackin (*δ* = +0.41, *p* = 0.032) trees relative to temperate terrestrial; north polar marine samples are more imbalanced under both Colless (*δ* = +0.52, *p* = 0.005) and Sackin (*δ* = +0.56, *p* = 0.003). The full set of latitudinal tree-shape contrasts is given in Table S5.

### Two-axis structure and the most divergent corner

Taken together, biome and latitude define two orthogonal axes of viral genomic divergence in the study. The biome axis is the larger and more consistent of the two: terrestrial viromes are systematically further from current IMG/VR references than marine ones, across every latitudinal band and across all three tree-shape statistics. The latitudinal axis modulates this baseline contrast: south polar samples sit at the high-divergence end of both biomes, with the terrestrial Antarctic stratum occupying the most extreme corner of the two-axis landscape. Both axes survive the move from per-distance Kruskal-Wallis tests (which are inflated by pseudoreplication of millions of within-tree pairwise distances) to per-SRA Wilcoxon tests at the proper unit of biological replication. Diagnostic regressions of per-SRA bridging edge against assembly N50 (Fig. S2) confirm that neither axis is driven by differential assembly contiguity in the cohort.

## Discussion

Our re-analysis of 120 environmental DNA metagenomes uncovers a structuring of viral genomic divergence that is dominantly biome-driven rather than latitude-driven. Terrestrial viromes are systematically further from IMG/VR reference clades than marine viromes, in every latitudinal band and across all three tree-shape statistics; the polar effects previously emphasized in the literature operate within and on top of this stronger biome effect, with the terrestrial Antarctic stratum occupying the most divergent corner of the resulting landscape. This reframing has both a methodological and a biological interpretation.

Methodologically, much of the apparent polar excess in earlier reports reflects the unit of statistical replication used. Pairwise patristic distances within a CoPHSe tree are not independent observations: variance partitioning of our 1.9 *×* 10^6^ per-CoPHSe values assigns 76.5% of variance to within-sample sources and only 23.5% to between-sample sources, so per-distance Kruskal-Wallis tests inflate degrees of freedom by approximately three orders of magnitude. Once this is corrected by aggregating to per-SRA medians, the polar-versus-equatorial bridging-edge contrast remains significant for the Antarctic in both biomes, but the within-clade MPD contrast does not reach *α* = 0.05 at the current *n*=16-24 per stratum. The marine-versus-terrestrial contrast survives the same correction more robustly because it operates on a larger axis (combining samples across all three regions, with *n* in the high 50s per biome) and reflects a larger effect size.

Biologically, the biome contrast is consistent with longstanding microbial-ecology observations. Soils are spatially structured at the millimeter scale, with pH, water potential, oxygen, and organic-matter gradients producing more independent ecological niches per unit volume than seawater [13]. Soil bacterial and archaeal communities are correspondingly more diverse, more dispersal-limited, and more endemic than their marine counterparts, and the viral component of these communities inherits this structure through tight host coupling [12, 14]. Soils additionally archive ancient microbial and viral lineages through permafrost preservation [15, 25]; this archival mode introduces a class of evolutionary distances rarely sampled in marine environments, and potentially explains why terrestrial Antarctic samples sit at the extreme corner of the two-axis landscape we report here. In contrast, ocean circulation, mesoscale eddies, and the relatively well-connected nature of pelagic environments homogenize marine viromes more efficiently and keep their lineages closer to a single recognizable backbone of reference sequences [11].

A reference-coverage component undoubtedly amplifies the biome contrast. IMG/VR v4.1 has been built from a viral sequencing record that historically over-represented marine ecosystems, most prominently through the Tara Oceans datasets, relative to soils, even though the soil-virome record has expanded substantially in recent years [12, 10]. Within-IMG/VR reference MPDs in our data differ subtly between biomes in a region-dependent way (Fig. S6); however, the magnitude of these reference-set differences is roughly five times smaller than the marine-versus-terrestrial bridging-edge contrast, so the biological signal cannot be fully reduced to differential reference coverage. A practical implication is that the highest-yield investments in expanding viral reference databases are now probably in terrestrial systems, and especially in terrestrial polar systems.

The polar effects we recover are smaller in magnitude than the biome effect but biologically meaningful. Antarctic samples in both biomes are significantly further from IMG/VR clades than equatorial samples; Arctic samples trend in the same direction without reaching significance at the current sample size. Three complementary mechanisms remain plausible for the polar amplification. Geographic isolation, particularly Antarctic circumpolar currents and atmospheric circulation [50], restricts gene flow and promotes lineage divergence through drift and host specialization; the elevated internal divergence and increased tree imbalance we observe in south polar samples are consistent with this mechanism, and recent Antarctic surveys similarly report high viral endemism [51]. Seasonal and environmental pulsing restructure microbial host communities, producing cyclical viral turnover [24]. Finally, climate-driven acceleration reshapes host ranges and selection regimes; the Arctic, warming nearly four times faster than the global mean [22], epitomizes conditions that favor rapid diversification. These polar processes act on top of the biome baseline and disproportionately affect terrestrial systems, where ancient viral reservoirs in permafrost [15] and active soil-microbiome turnover under thaw [28, 52] together exposes a class of divergence not represented in current marine-dominated reference collections.

Several caveats temper the interpretation. Within-clade MPDs do not reach significance (*α* = 0.05) at sample-level resolution despite moderate effect sizes (*δ* = 0.19-0.29 for south polar versus equatorial contrasts), reflecting limited sample numbers per stratum rather than the absence of biological signal; expanded sampling will be required for formal statistical confirmation. Following QC filtering, most north polar marine datasets originated from only two BioProjects representing a narrow spatial transect near Alaska (Fig. 2; Table S2), so north polar marine divergence estimates may partly reflect localized ecological structure. The bridging-edge metric, which retains statistical significance for both biome and polar contrasts, is jointly shaped by the diversity of recruited reference sequences in IMG/VR and by genuine evolutionary distance (see Fig. S7) and shown that its magnitude is substantially smaller than the biological contrast it cannot explain. Diagnostic regressions against assembly N50 (Fig. S2) confirm that neither the biome nor the polar contrast is driven by differential assembly contiguity.

The two-axis structure we describe has direct implications for biodiversity baselines and predictive ecology. As polar ecosystems experience accelerating climate change [22, 52], permafrost thaw and shifting hydrological regimes will preferentially mobilize viral reservoirs from the very corner of our landscape (terrestrial Antarctic and Arctic) that is most distant from current references. Strategic sampling and database expansion should therefore prioritize terrestrial polar systems rather than continuing to add depth where coverage is already dense. Adherence to the MIUViG framework [14] will ensure that the contextual metadata needed for biome-aware analyses (collection date, geographic coordinates, environmental context) accompany newly recovered genomes. Beyond polar contexts, the CoPHSe framework previously introduced [21] and exploited here, together with the sample-level inferential ladder and tree-shape diagnostics added in this work, offers a standardized, reproducible approach to compare biomes and latitudes on a common footing.

## Acknowledgements

We would like to thank the Digital Research Alliance of Canada for providing us with compute time.

## Author contributions

S.A.B. conceptualized the study. V.K. collected the data, performed the analyses, wrote the manuscript and designed all figures which were reviewed and edited by all authors.

## Data availability

All raw sequencing reads analyzed in this study are publicly available in NCBI’s Sequence Read Archive (https://www.ncbi.nlm.nih.gov/sra). All analysis scripts used in this study are available on GitHub at https://github.com/vkulk094/viral_phylo, with additional reproducibility scripts at https://github.com/sarisbro/data. Revision-stage scripts (sample-level inference, tree-shape pipeline, consolidated per-CoPHSe summary, biome-contrast tests) are included in the same repository under revision scripts/.

