## Supplementary material for "Terrestrial DNA viromes are systematically more divergent from reference databases than marine ones, with polar ecosystems amplifying the contrast": SI figures and tables

### Supplementary Figures

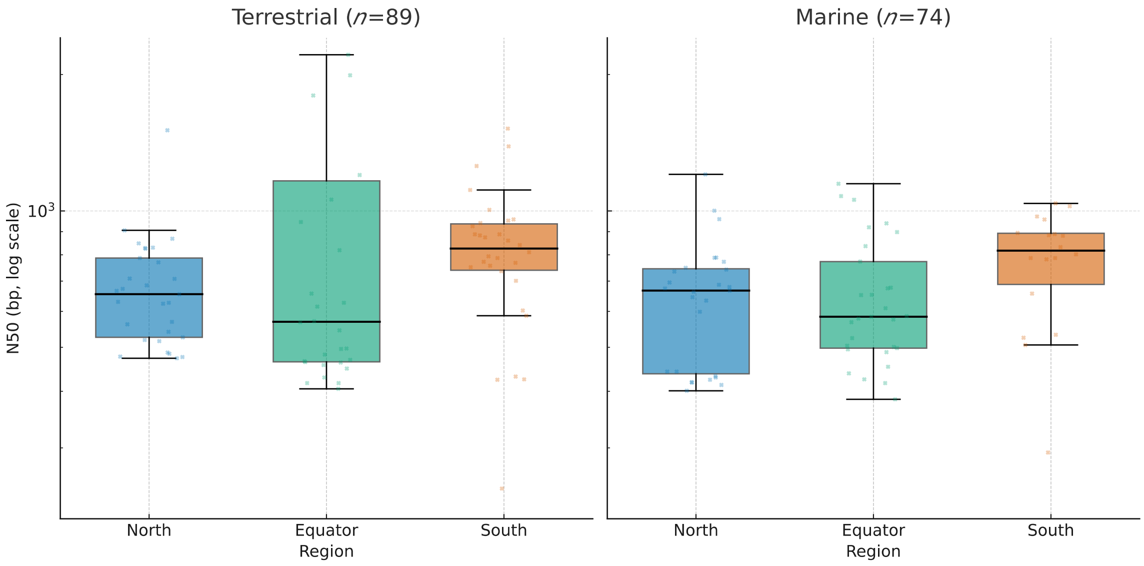


**Figure S1.** Box-plots of contig N50 values (log scale) by region within terrestrial and marine environments. Regions are ordered north (blue), temperate/equator (green), and south (vermillion). Kruskal-Wallis tests indicated no significant differences at $\alpha = 0.01$ (Terrestrial: $H = 5.12$, $p = 0.077$; Marine: $H = 6.54$, $p = 0.038$).

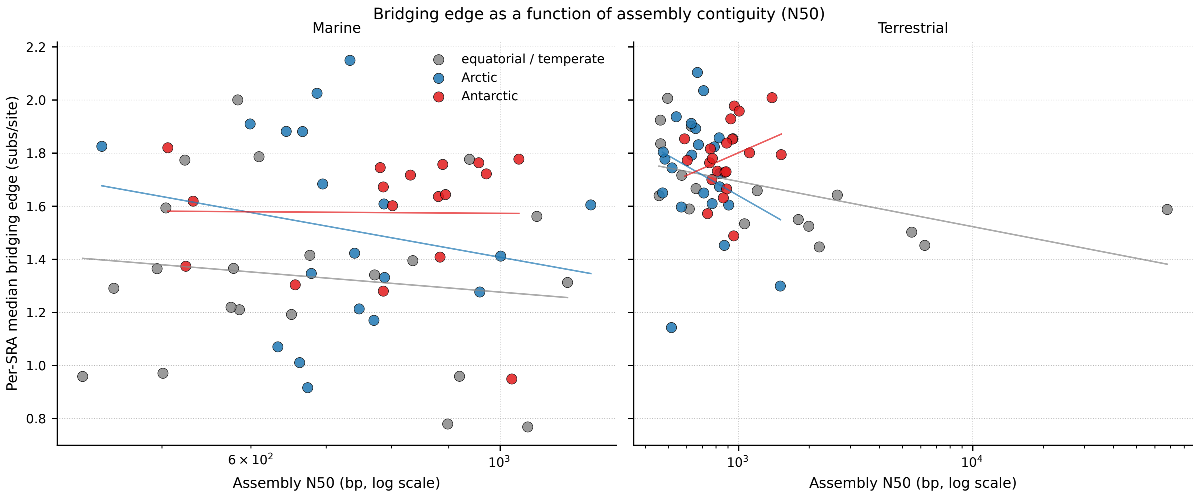


**Figure S2.** Assembly contiguity does not drive the divergence patterns we report. Per-SRA scatter of the median bridging-edge value (y-axis) against the assembly N50 in base pairs (x-axis, log scale), separated by biome and colored by latitudinal band. Per-region log-linear smoothers are shown. Within each biome, both the biome and the latitudinal patterns reported in main-text Fig. 6 and Supplementary Table S2 persist across the full range of N50 values represented in the cohort, indicating that the divergence patterns are not artifacts of differential assembly contiguity (which differs across marine strata at $p = 0.038$).

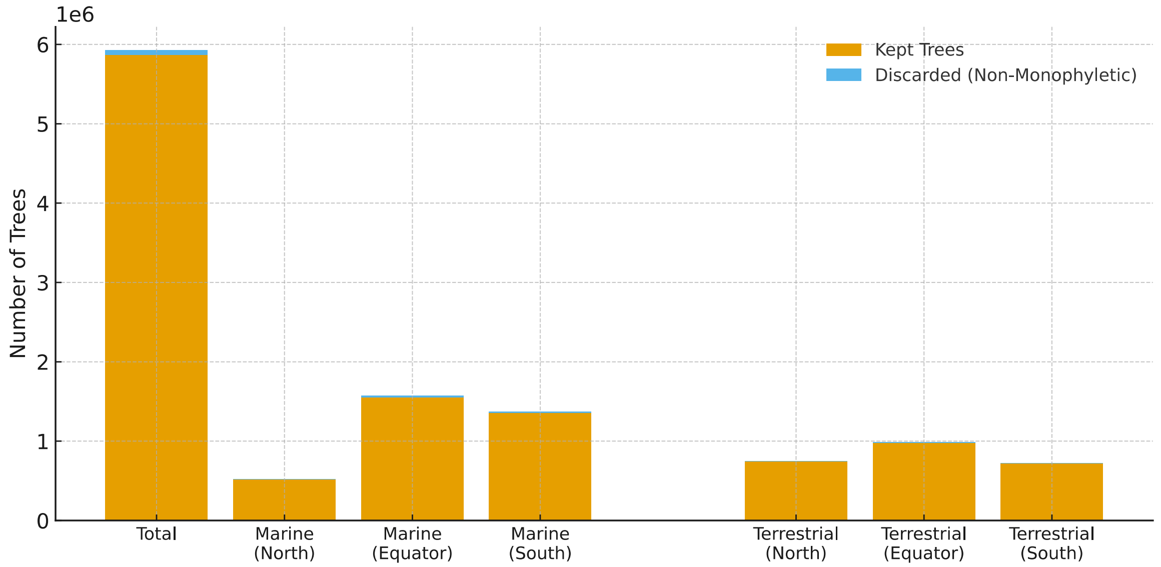


**Figure S3.** Proportion of phylogenetic trees retained versus discarded due to non-monophyly across regions. Bars show the total number of valid trees (orange) and the fraction discarded because they were non-monophyletic (blue). Results are shown for the global total as well as subdivided into Marine (north, temperate/equator, south) and Terrestrial (north, temperate/equator, south) regions. Across all regions, discarded trees represent only a very small fraction of the total (0.35%).

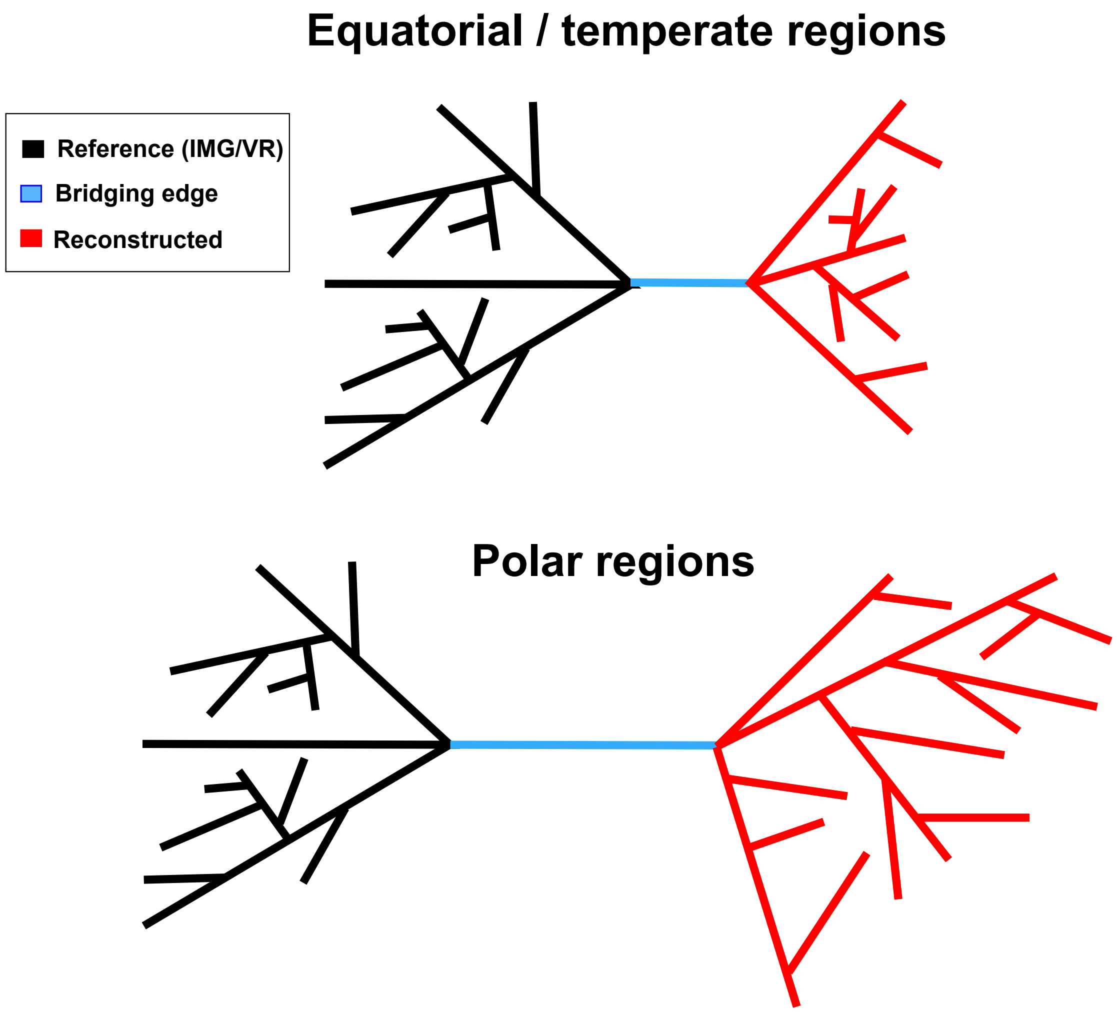


**Figure S4.** Schematic representation of the expected branch length patterns of phylogenetic trees in equatorial/temperate regions compared to polar regions.

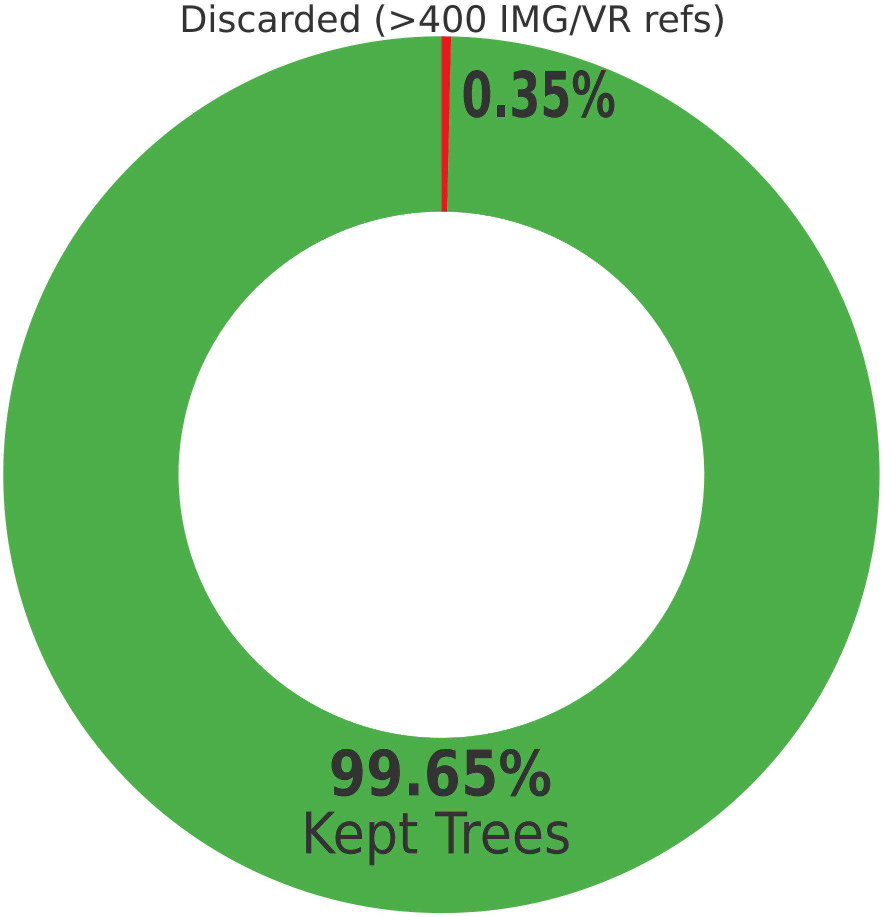


**Figure S5.** Global proportion of phylogenetic trees excluded for exceeding 400 IMG/VR references. The 400-reference threshold was applied uniformly to control the quadratic scaling of patristic distance calculations in R, which become disproportionately ($\mathcal{O}(n^2)$) expensive in memory and runtime as the number of references increases. The majority (99.65%) trees were kept and only a fraction (0.35%) were discarded.

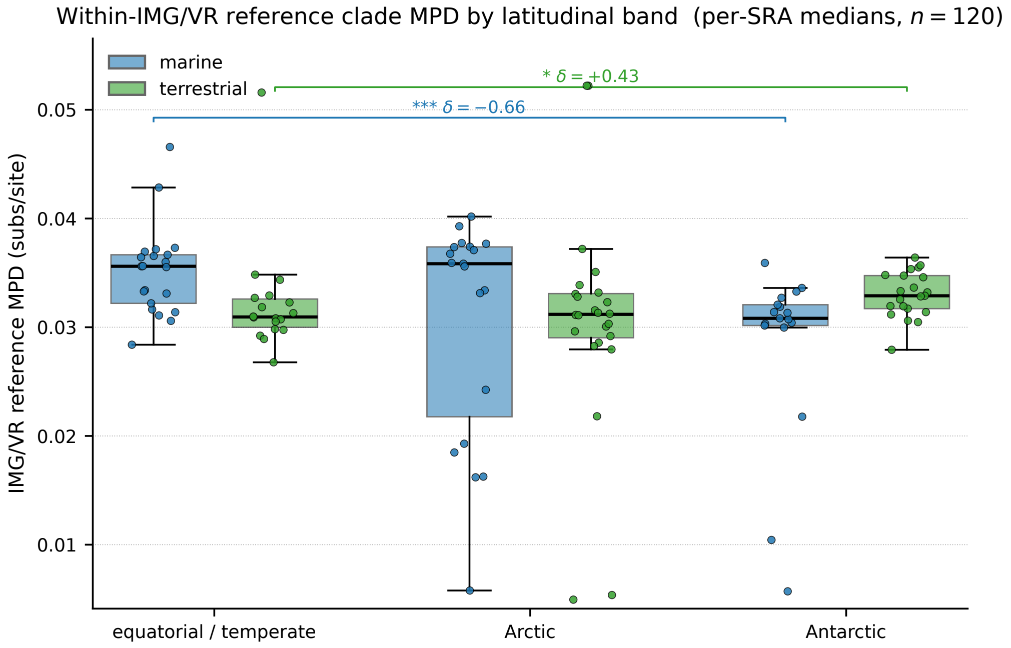


**Figure S6.** Within-IMG/VR reference clade MPD per SRA across latitudinal bands. Boxplots and per-SRA jittered points for the mean pairwise patristic distance (MPD) between IMG/VR reference sequences within each CoPHSe, aggregated to per-SRA medians ($n = 120$ samples). Marine south polar samples recruit references that are tighter on average (Cliff's $\delta = -0.66$, $p < 0.001$), while terrestrial south samples recruit references that are slightly looser ($\delta = +0.43$, $p = 0.023$). These asymmetries reflect the latitudinal composition of the IMG/VR collection itself and are an important caveat for the interpretation of absolute bridging-edge values; their magnitude is, however, roughly five times smaller than the marine-versus-terrestrial bridging-edge contrast reported in main-text Fig. 6.

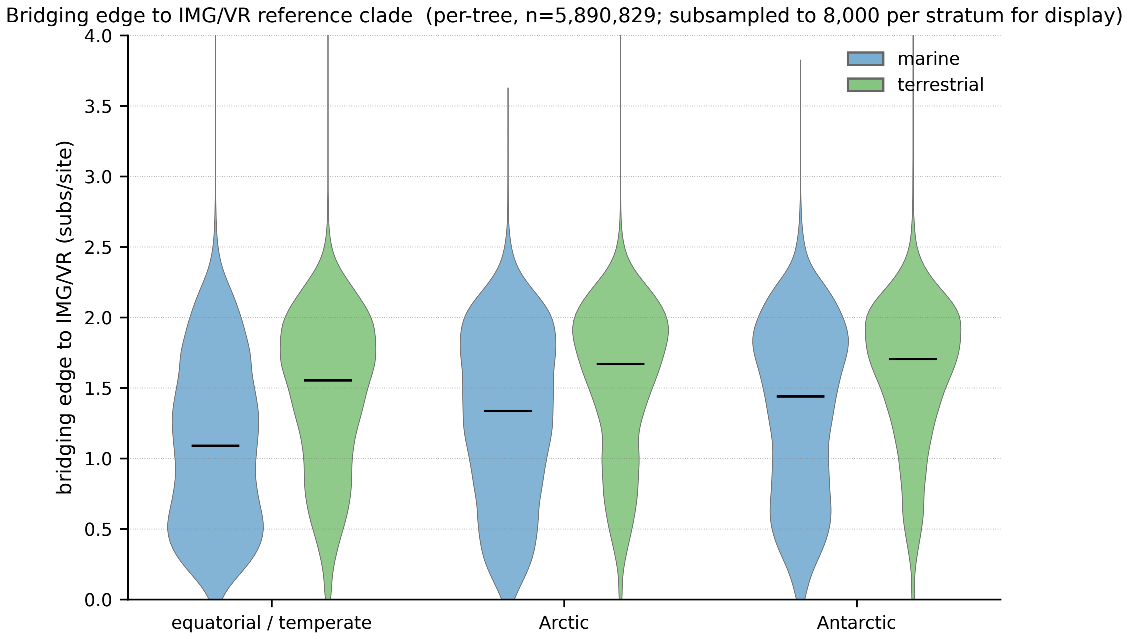


**Figure S7.** Per-tree distributions of the bridging-edge metric across latitudinal bands. Violin plots show the full distribution of bridging-edge values across all 5,890,829 trees with a defined bridging edge. For display, each stratum was subsampled to 8,000 trees. The polar shift toward greater bridging edges is visible as a rightward shift of the bulk of the distribution in both polar biomes; the corresponding sample-level Wilcoxon test on per-SRA medians is reported in Supplementary Table S1 and visualized at sample-level resolution in main-text Fig. 6.

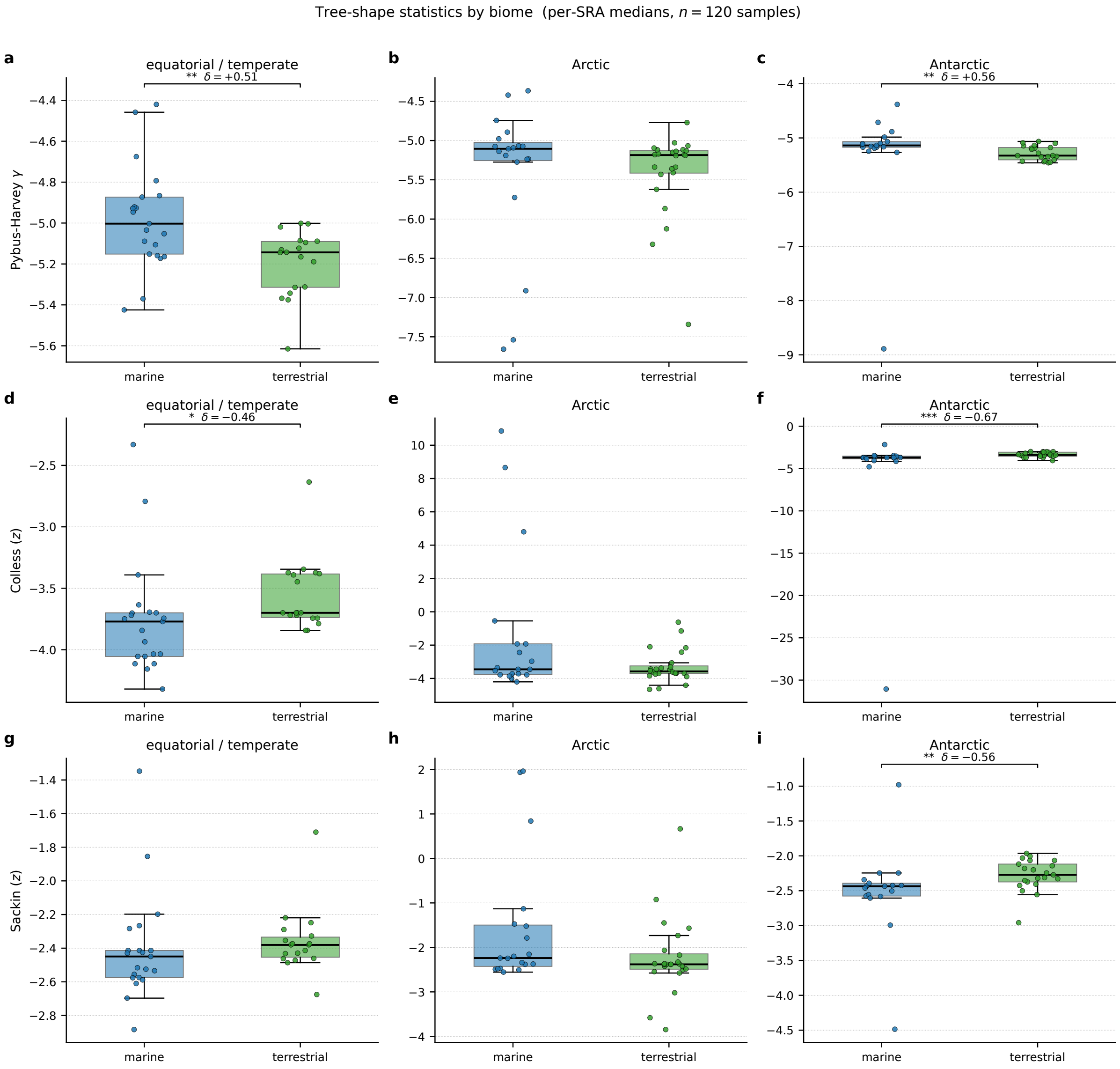


**Figure S8.** Tree-shape statistics by biome across latitudinal bands. Boxplots and per-SRA jittered points for (a-c) Pybus-Harvey $\gamma$, (d-f) Yule-normalized Colless imbalance, and (g-i) Yule-normalized Sackin imbalance, with biome on the x-axis and one column per latitudinal band. Significance brackets show sample-level Wilcoxon rank-sum contrasts between marine and terrestrial within each band, with Cliff's $\delta$ effect sizes inline (* $p < 0.05$, ** $p < 0.01$, *** $p < 0.001$); only contrasts with $p < 0.05$ are annotated. Terrestrial trees are significantly more imbalanced and more deeply branching than marine trees in the equatorial and Antarctic bands, with the Antarctic biome contrast reaching significance on all three tree-shape statistics. Arctic contrasts trend in the same direction without reaching $\alpha = 0.05$. The full 24-row contrast table is given in Supplementary Table S4.

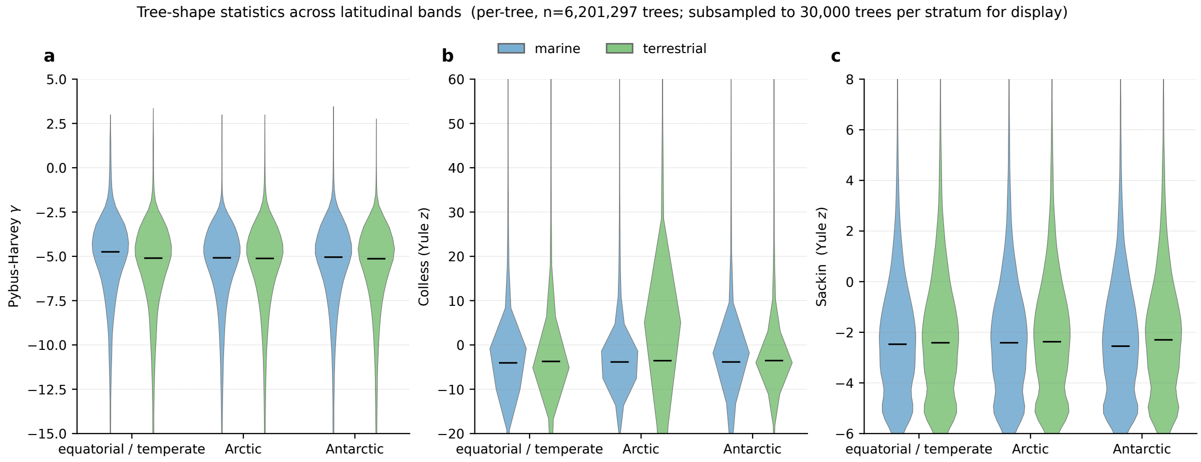


**Figure S9.** Per-tree distributions of tree-shape statistics across latitudinal bands. Violin plots show the full distribution of (a) Pybus-Harvey $\gamma$, (b) Yule-normalized Colless imbalance, and (c) Yule-normalized Sackin imbalance across all 6,201,297 CoPHSe trees with $\geq 6$ environmental tips. For display, each stratum was subsampled to $3 \times 10^4$ trees. The corresponding sample-level Wilcoxon tests on per-SRA medians are summarized in Supplementary Table S5.

### Supplementary Tables S1 and S2 (separate Excel file).

Supplementary Tables S1 and S2 are provided as a separate Excel file (`Kulkarni.Aris-Brosou_SI2-DNAT` [illegible] Table S1 lists the full SRA accession identifiers for all 180 datasets initially retrieved and the 120 that passed quality control, together with BioProject identifiers, run-level metadata, and study titles. Table S2 lists the BioSample geographic coordinates (latitude, longitude), collection dates, environmental metadata (biome, environment material, depth where applicable), and the geographic feature that motivated each sample's inclusion in the cohort.

### Supplementary Table S3. Sample-level Wilcoxon contrasts on divergence metrics, latitudinal axis.

**Table S3.** Sample-level Wilcoxon rank-sum tests (one observation per SRA, $n = 16$-24 per stratum) on per-CoPHSe medians of the within-clade reconstructed MPD (`gene_mpd`), the bridging edge to the IMG/VR reference clade (`bridging_edge`), and the within-IMG/VR reference MPD (`imgvr_mpd`), comparing each polar band to the equatorial / temperate baseline of the same biome. Bold rows indicate $p < 0.05$. Cliff's $\delta$ effect sizes are reported; 95% CIs from 2,000 bootstrap replicates on the difference in medians.

| metric | biome | contrast | Δ median | 95% CI | Cliff $\delta$ | $p$ |
|---|---|---|---|---|---|---|
| gene_mpd | marine | north vs eq | −0.184 | [−0.751, +0.265] | −0.244 | 0.220 |
| gene_mpd | marine | south vs eq | +0.494 | [−0.066, +0.749] | +0.288 | 0.139 |
| gene_mpd | terrestrial | north vs eq | +0.086 | [−0.526, +0.407] | +0.031 | 0.883 |
| gene_mpd | terrestrial | south vs eq | +0.284 | [−0.249, +0.494] | +0.188 | 0.337 |
| bridging_edge | marine | north vs eq | +0.082 | [−0.100, +0.513] | +0.278 | 0.136 |
| **bridging_edge** | **marine** | **south vs eq** | **+0.303** | **[+0.067, +0.498]** | **+0.429** | **0.026** |
| bridging_edge | terrestrial | north vs eq | +0.144 | [−0.020, +0.259] | +0.310 | 0.091 |
| **bridging_edge** | **terrestrial** | **south vs eq** | **+0.139** | **[+0.018, +0.252]** | **+0.413** | **0.029** |
| imgvr_mpd | marine | north vs eq | 0.000 | [−0.011, +0.004] | −0.008 | 0.978 |
| **imgvr_mpd** | **marine** | **south vs eq** | **−0.005** | **[−0.006, −0.001]** | **−0.664** | **0.001** |
| imgvr_mpd | terrestrial | north vs eq | 0.000 | [−0.002, +0.002] | −0.028 | 0.889 |
| **imgvr_mpd** | **terrestrial** | **south vs eq** | **+0.002** | **[+0.000, +0.004]** | **+0.429** | **0.023** |

The intraclass correlation coefficient on per-CoPHSe `gene_mpd` is 0.235: only 23.5% of variance lies between SRAs, with the remaining 76.5% within. The original $\chi^2$ values shown in main-text Figs. 3 and 4 reflect treating $\sim 10^6$ within-tree pairwise distances as independent observations and inflate degrees of freedom by approximately three orders of magnitude. The contrasts reported here use one observation per SRA, the unit of biological replication.

#### Supplementary Table S4. Sample-level Wilcoxon contrasts, biome axis (marine vs terrestrial).

The biome contrast on the bridging-edge metric is statistically significant in every latitudinal band, with consistent directionality (terrestrial > marine) and moderate-to-large effect sizes ($|\delta| = 0.38$-$0.63$). This is the larger of the two structuring axes in the cohort, and motivates the biome-led framing of the main manuscript. Within the Antarctic band, all three tree-shape statistics also show significant biome contrasts (Cliff's $\delta$ between $-0.56$ and $-0.67$), with terrestrial trees more imbalanced and more deeply branching than marine trees.

**Table S4.** Sample-level Wilcoxon rank-sum tests (one observation per SRA) comparing marine versus terrestrial biomes on all six revision metrics. "ALL" rows pool across the three latitudinal bands; per-region rows test within each band. Effect sizes are Cliff's $\delta$ (negative values indicate marine < terrestrial); 95% CIs from 2,000 bootstrap replicates on the difference in medians. Bold rows indicate $p < 0.05$.

| metric | region | $n_M : n_T$ | $\Delta$ median (M−T) | Cliff $\delta$ | 95% CI | $p$ |
|---|---|---|---|---|---|---|
| gene_mpd | ALL | 53 : 58 | −0.226 | −0.122 | [−0.519, +0.128] | 0.271 |
| gene_mpd | eq | 20 : 17 | −0.310 | −0.135 | [−0.543, +0.214] | 0.493 |
| gene_mpd | north | 16 : 21 | −0.579 | −0.226 | [−0.928, +0.226] | 0.250 |
| gene_mpd | south | 17 : 20 | −0.099 | −0.047 | [−0.321, +0.454] | 0.819 |
| **bridging_edge** | **ALL** | **57 : 63** | **−0.329** | **−0.497** | **[−0.421, −0.105]** | **$< 10^{-5}$** |
| **bridging_edge** | **eq** | **21 : 18** | **−0.300** | **−0.630** | **[−0.472, −0.159]** | **$8 \times 10^{-4}$** |
| **bridging_edge** | **north** | **19 : 24** | **−0.362** | **−0.377** | **[−0.521, +0.034]** | **0.037** |
| **bridging_edge** | **south** | **17 : 21** | **−0.137** | **−0.591** | **[−0.372, −0.015]** | **0.002** |
| imgvr_mpd | ALL | 57 : 63 | +0.002 | +0.204 | [−0.000, +0.004] | 0.055 |
| **imgvr_mpd** | **eq** | **21 : 18** | **+0.005** | **+0.587** | **[+0.001, +0.006]** | **0.002** |
| imgvr_mpd | north | 19 : 24 | +0.005 | +0.259 | [−0.007, +0.007] | 0.153 |
| **imgvr_mpd** | **south** | **17 : 21** | **−0.002** | **−0.518** | **[−0.004, −0.000]** | **0.007** |
| $\gamma$ | **ALL** | **57 : 63** | **+0.089** | **+0.431** | **[+0.033, +0.245]** | **$< 10^{-4}$** |
| $\gamma$ | **eq** | **21 : 18** | **+0.140** | **+0.513** | **[+0.017, +0.322]** | **0.007** |
| $\gamma$ | north | 19 : 24 | +0.080 | +0.246 | [−0.059, +0.271] | 0.175 |
| $\gamma$ | **south** | **17 : 21** | **+0.188** | **+0.557** | **[+0.025, +0.267]** | **0.004** |
| **Colless ($z$)** | **ALL** | **57 : 63** | **−0.265** | **−0.272** | **[−0.329, −0.009]** | **0.010** |
| **Colless ($z$)** | **eq** | **21 : 18** | **−0.071** | **−0.458** | **[−0.489, +0.000]** | **0.015** |
| Colless ($z$) | north | 19 : 24 | +0.125 | +0.173 | [−0.265, +1.610] | 0.340 |
| **Colless ($z$)** | **south** | **17 : 21** | **−0.312** | **−0.672** | **[−0.637, −0.164]** | **$4 \times 10^{-4}$** |
| Sackin ($z$) | ALL | 57 : 63 | −0.052 | −0.166 | [−0.119, +0.000] | 0.117 |
| Sackin ($z$) | eq | 21 : 18 | −0.069 | −0.307 | [−0.182, +0.012] | 0.105 |
| Sackin ($z$) | north | 19 : 24 | +0.142 | +0.292 | [−0.015, +0.857] | 0.106 |
| **Sackin ($z$)** | **south** | **17 : 21** | **−0.163** | **−0.557** | **[−0.363, −0.071]** | **0.004** |

### Supplementary Table S5. Sample-level tree-shape contrasts, latitudinal axis.

**Table S5.** Sample-level Wilcoxon rank-sum tests on per-SRA medians of three tree-shape statistics computed across 6,201,297 CoPHSe trees with $\geq 6$ environmental tips (all 120 samples). Effect sizes are Cliff's $\delta$; bootstrap 95% confidence intervals on the difference in medians. Bold rows indicate $p < 0.05$.

| metric | biome | contrast | $\Delta$ median | 95% CI | Cliff $\delta$ | $p$ |
|---|---|---|---|---|---|---|
| $\gamma$ | marine | north vs eq | −0.102 | [−0.309, +0.030] | −0.313 | 0.093 |
| $\gamma$ | marine | south vs eq | −0.135 | [−0.245, −0.004] | −0.333 | 0.083 |
| $\gamma$ | terrestrial | north vs eq | −0.042 | [−0.237, +0.121] | −0.269 | 0.144 |
| $\gamma$ | **terrestrial** | **south vs eq** | **−0.183** | **[−0.239, +0.032]** | **−0.397** | **0.036** |
| **Colless ($z$)** | **marine** | **north vs eq** | **+0.317** | **[+0.024, +1.845]** | **+0.519** | **0.005** |
| Colless ($z$) | marine | south vs eq | +0.067 | [−0.072, +0.341] | +0.188 | 0.332 |
| Colless ($z$) | terrestrial | north vs eq | +0.122 | [−0.239, +0.314] | +0.155 | 0.401 |
| **Colless ($z$)** | **terrestrial** | **south vs eq** | **+0.309** | **[−0.010, +0.533]** | **+0.466** | **0.014** |
| **Sackin ($z$)** | **marine** | **north vs eq** | **+0.210** | **[+0.045, +0.928]** | **+0.561** | **0.003** |
| Sackin ($z$) | marine | south vs eq | +0.014 | [−0.142, +0.130] | −0.025 | 0.907 |
| Sackin ($z$) | terrestrial | north vs eq | 0.000 | [−0.060, +0.094] | +0.039 | 0.839 |
| **Sackin ($z$)** | **terrestrial** | **south vs eq** | **+0.108** | **[+0.015, +0.263]** | **+0.405** | **0.032** |
